# Novel single molecule imaging approaches reveal structure-function alterations to the nuclear pore complex in early *C9ORF72*-associated TDP-43 proteinopathy

**DOI:** 10.64898/2026.02.17.706337

**Authors:** Seoungjun Lee, Sarah Mizielinska

**Author notes:** Joint corresponding authors.

## Abstract

Nucleocytoplasmic transport through the nuclear pore complex is essential for the maintenance of cellular homeostasis by regulating the movement of molecules between the nucleus and the cytoplasm. This process becomes dysfunctional in many diseases but has been particularly implicated in amyotrophic lateral sclerosis (ALS) and frontotemporal dementia (FTD) linked to the *C9ORF72* mutation and associated aberrantly produced polypeptides. To directly study nucleocytoplasmic transport in intact non-genetically modified cells we have developed two new approaches for single molecule tracking through nuclear pore subdomains and super-resolved imaging of nuclear pore structural organisation. Using these techniques we have examined the early impact of the neurotoxic *C9ORF72* polypeptide poly(glycine-arginine) on nuclear pore transport dynamics and structure. We find that soluble poly(glycine-arginine) peptides can disrupt molecular flow of passive cargo predominantly during nuclear export which is associated with altered structural organisation and molecular interaction of nuclear basket and central nuclear pore complex domains. These changes converge with perturbed nucleocytoplasmic homeostasis of the key ALS/FTD pathological protein TDP-43 and begin to explain this initiating step in disease.

---

Nucleocytoplasmic transport is a fundamental process critical for gene expression, allowing the movement of RNAs and pre-ribosomal complexes, but also for the responsiveness of cells to stimuli or stress via the movement of DNA/RNA binding proteins that regulate gene expression ^1, 2^. Virtually all nucleocytoplasmic transport occurs via large macromolecular nuclear pore complexes (NPCs) which generate a channel that spans the nuclear envelope comprised of multiple copies of proteins called nucleoporins organised into structural and flexible domains ^3^. The ability of molecules to pass through the NPC is determined by their biophysical properties, including size, surface charge and hydrophobicity ^4, 5^. However, in general relatively inert molecules smaller than approximately 40 kDa can rapidly diffuse through NPCs, while larger and more charged molecules require specific transport receptors for effective transport ^3, 4, 6^. Thus, nucleocytoplasmic transport through the NPC is tightly controlled, and interference with either its structure or function has significant consequences for cellular homeostasis and resilience in disease ^7^.

A major pathological hallmark of the overlapping neurodegenerative disorders amyotrophic lateral sclerosis (ALS) and frontotemporal dementia (FTD) is the altered homeostasis of RNA binding proteins with displacement from the nucleus to the cytoplasm and subsequent aggregation ^8-10^. Indeed, nucleocytoplasmic mislocalisation of the RNA binding protein TDP-43 occurs in 97 percent of ALS cases and almost half of FTD cases, in addition to being present in a range of other neurological disorders and is highly associated with regions of neurodegeneration and in Alzheimer’s disease, cognitive impairment ^11-14^. The most common genetic cause in ALS/FTD is a non-coding repeat expansion mutation in the *C9ORF72* gene ^15-17^. In addition to partial loss of protein function and repeat RNA toxicity, the repeat expansion can be translated into aberrant dipeptide repeat (DPR) polypeptides ^18^, including the particularly toxic polypeptide poly-(glycine-arginine) (polyGR) ^19-23^. This arginine rich polypeptide has been shown to bind to low complexity domain containing proteins including nuclear pore proteins ^24-26^ and associate with TDP-43 pathology ^27-32^. However, the precise mechanisms linking polyGR, nucleocytoplasmic transport dynamics and TDP-43 mislocalisation are yet to be determined due to a lack of appropriate means to study this process.

To overcome the technical constraints that have limited our understanding of nucleocytoplasmic transport, we established novel integrative methodologies that leverage the complementary strengths of super-resolution imaging and deep learning–based analysis. These tools enable direct visualization and quantification of single molecule nucleocytoplasmic transport dynamics at sub-domain resolution and nuclear pore architecture in intact, unmodified cells, offering a transformative advance for dissecting transport mechanisms with molecular precision. Using these approaches, we demonstrate the impact of the aberrant polypeptide polyGR and the association between altered molecular flow and structural organisation of NPCs and early TDP-43 pathology in *C9ORF72* ALS/FTD.

## Results

### Defining nuclear pore sub-compartment nucleocytoplasmic transport dynamics in an intact unmodified cellular system

NPCs are comprised of smaller complexes organised predominantly into ring-based structures but can also be denoted as having three main domains – the nuclear basket, central domain (including the pore channel itself) and cytoplasmic filaments ^3^. In order to directly study nucleocytoplasmic transport, single molecule tracking must be performed. This overcomes the limitations of understanding from common summative studies in which nuclear and cytoplasmic levels are assessed, and contributions of protein production, turnover and transport cannot be segregated, or, where transport is inferred from altered compartment accumulation upon selective transport receptor inhibition where relevant inhibitors exist. From the first reports, studies predominantly utilised permeabilised cells or isolated nuclei to perform single molecule nucleocytoplasmic tracking in order to permit the controlled application of exogenous cargo in an artificial cytoplasm buffer ^33-35^. However, cell permeabilization can alter nuclear pore composition and permeability ^36, 37^. In addition, cytoplasmic homeostasis can also play key roles in modulating nucleocytoplasmic transport, particularly in neurodegenerative disease ^38-40^. Thus, we aimed to overcome these limitations by developing a novel methodology to monitor single molecule nucleocytoplasmic transport in intact cells using a combination of super-resolution microscopy, single molecule tracking and deep learning (Fig. 1).

**Fig. 1.**
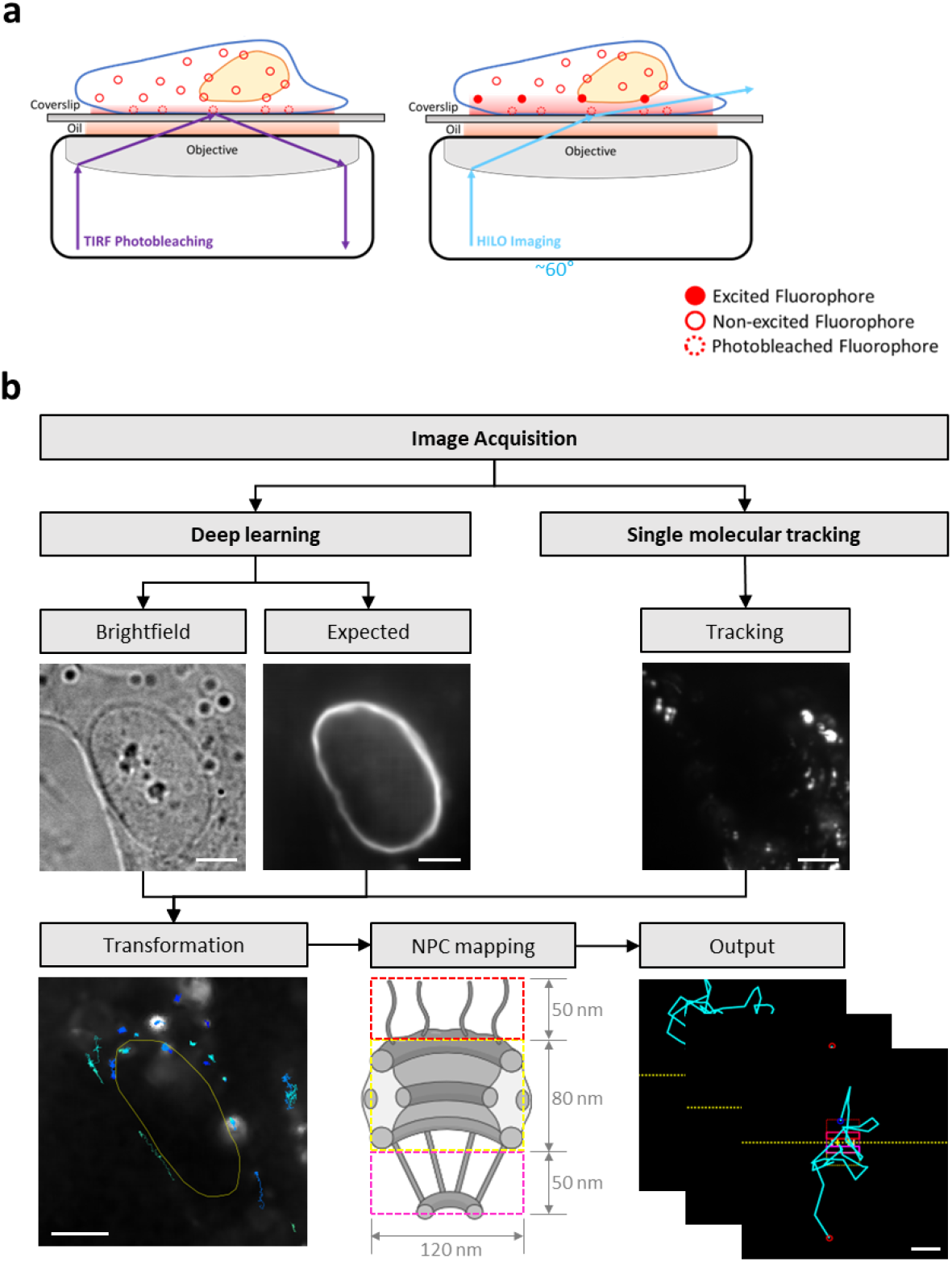
Overview for single molecule tracking of individual nuclear pore subdomain dynamics. Process involves a) the single molecular imaging technique of TIRF (Total Internal Reflection Fluorescence) bleaching of the sample close to the coverslip (<1 µm) and HILO (Highly Inclined and Laminated Optical sheet) imaging to excite single molecules of cargo 2 µm above the coverslip at the lower edge of the nuclear envelope in intact cells using 10,000 images, 53 frames per second, for 3 minutes. b) Single molecule image acquisition and tracking is transformed in combination with deep learning to predict the nuclear envelope position using brightfield imaging training using nuclear envelope marker mCherry-lamin B1 (expected) and mapping to a nuclear pore complex (NPC) domain model to output single molecule tracks through individual nuclear pore complexes in intact non-genetically modified cells. Scale bars for brightfield, expected, tracking, and transformation are 5 μm, and for output 100 nm.

One of the major challenges of single molecule tracking across the NPC is the depth at which the nuclear envelope is situated within live intact cells. We used the neuroblastoma cell line SH-SY5Y and incubated cells with a small inert CF640R-tagged 10 kDa dextran cargo which could permeate the cell plasma membrane. This cargo represents a small inert cargo and thus transport receptor-independent (commonly known as passive) nucleocytoplasmic transport. As single molecule tracking using a standard TIRF (total internal reflection fluorescence) approach is not suitable for the sample depth required, we used HILO (highly inclined and laminated optical sheet) imaging, which uses a low-powered laser at below-critical angle to increase illumination of sample further above the coverslip, while maintaining selectivity of single molecules. This was optimised to 1.67° below the critical angle, giving a final angle of 59.56°, for acquisition at a depth of approximately 2 µm above the coverslip and at the lower edge of the nuclear envelope, with some adjustment for focus by further reducing the incident laser beam angle. To reduce noise, bleaching close to the coverslip was also performed using a high-powered laser at an angle of ∼68° for 1 minute. For sufficient sampling of single molecule tracking through the NPC, HILO imaging was performed for 10,000 frames at 53 frames per second (∼ 3 minutes, Fig. 1a, Extended Data Fig. 1).

Single molecule images were denoised using CANDLE (Collaborative Approach for Enhanced Denoising under Low-light Excitation ^41^, Supplementary Video 1), which was designed for use in low signal-to-noise ratio conditions typical of deep image acquisition in biological samples. Custom analysis code was used for single molecule spot detection and tracks determined using the nearest neighbour method. To avoid the requirement for cells genetically modified for labelling of a nuclear envelope or nuclear pore component or co-application of a cargo known to accumulate in NPCs, which may all alter transport, nuclear envelope position was determined using a deep learning approach on brightfield images collected simultaneously. Training on ∼10,000 images was performed using samples acquired for both brightfield and the nuclear envelope marker mCherry-lamin B1, with an outward adjustment applied to define the nuclear envelope central position, accounting for lamin B1 localisation within the lamina at the inner nuclear membrane interface (Extended Data Fig. 2). Single molecule tracks that did not intersect with the central location of the nuclear envelope, as determined by deep learning ^42^, were excluded. As the nuclear envelope provides a significant barrier to nucleocytoplasmic transport, transversal across the NPC will be the principal route of transport of our small inert cargo. Thus, after rotation in line with the nuclear envelope, single molecule tracks were overlaid on an NPC domain model of known mammalian NPC dimensions ^3, 43^. To ensure robust analysis, we excluded tracks with overlapping localisations in space (<160 nm) or time (<1 s). Moreover, we stringently defined track interaction with the NPC (and not low affinity interactions with the nuclear envelope), only including tracks with ≥10 molecular positions aligned with NPC dimensions and ≥3 positions with the NPC central pore domain. Together, this image processing defined single molecule transport through single NPCs and NPC subdomains (Fig. 1b, Supplementary Videos 2 and 3).

**Figure 2.**
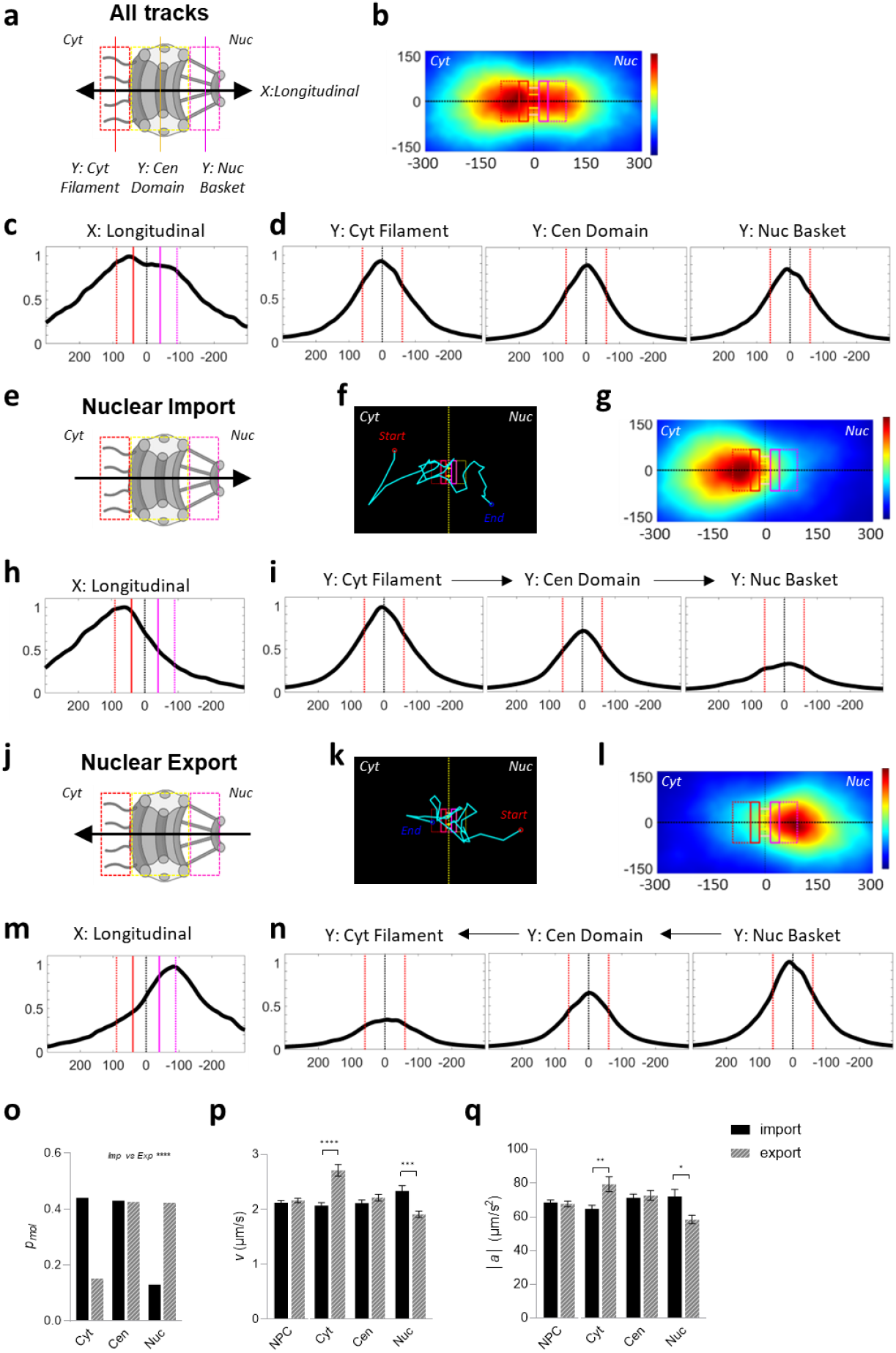
Single molecule passive nucleocytoplasmic transport dynamics defined by direction and subdomain. Neuroblastoma cell line SH-SY5Y was incubated with a small inert CF640R-tagged 10 kDa dextran cargo and single molecule tracking performed on intact cells. Dissection of tracks and dynamics: a-d, All tracks; e-i, nuclear import; j-n, nuclear export. Each dataset is presented with a nuclear pore complex (NPC) schematic with direction of transport overlaid with subdomain dimensions with dashed lines highlighting the cytoplasmic filament (red), central (yellow) and nuclear basket (pink) subdomains, and lines denoting longitudinal and transverse (across subdomain) line scan positions. Each data set includes (axis in nm) density heatmaps for molecular positions within tracks with overlaid NPC dimensions; an example track (cyan) on aligned nuclear envelope (yellow dashed line) with nuclear pore overlay and start (red) and end (blue) positions; comparative longitudinal (y = 0 in respect to central position on NPC) density profile through the NPC; transverse profiles through the midpoint of each of the cytoplasmic filament (x = −90nm), central (x = 0) and nuclear basket (x = +90nm) subdomains. Quantitative analysis of nuclear import (solid bars) versus nuclear export (diagonal striped bars) display data for o, proportional molecular density between different subdomains (pmol), p, velocity (v) and q, absolute acceleration (|a|) for total NPC or subdomains. Statistical analyses of molecular data: proportional molecular density – chi squared, velocity and absolute acceleration - one-way ANOVA with Dunnett’s multiple comparison on log-transformed data for total NPC, 2-way ANOVA with Dunnett’s multiple comparison vs control on log-transformed data for subdomains; *p<0.05, **p<0.01, ***p<0.001, ****p<0.0001; N>1000 for total NPC, N>145 for subdomains; raw numerical data in Source Data File for Fig. 2/3.

Final molecular tracks representing nucleocytoplasmic transport through NPCs (Fig. 2a) exhibit the expected higher density of interactions within the NPC boundary area, with a narrowing of molecular density as transport occurs through the central transport channel (Fig. 2b). This is also demonstrated by intensity line scans along the longitudinal axis of the nuclear pore (Fig. 2c). As expected for molecules moving through a cylindrical pore, transverse density plots through the cytoplasmic filaments, central and nuclear basket domains show a peak at the central position within all domains (Fig. 2d). Tracks were subsequently segregated into directionality to determine nuclear import (cytoplasmic starting point, Fig. 2e,f) and nuclear export (nuclear starting point, Fig. 2j,k). When separated, there is strong definition in density plots between the directions, both with teardrop shaped intensity boundaries, with nuclear import displaying higher molecular density in the cytoplasmic filament domain constricting to the central channel and then signal dispersal in the nuclear basket domain and nucleus (Fig. 2g), and the reverse with nuclear export (Fig. 2l). Again, this was corroborated by longitudinal intensity line scans with skewed distributions (Fig. 2h,m) and the variable height in transverse scans (Fig. 2i,n) across the respective subdomains. On quantification, 42-44% of molecular density was in each of the cytoplasmic filament and central channel domains for nuclear import with the remaining ∼13% in the nuclear basket, and for export ∼42% in nuclear basket and central domains and ∼15% in the cytoplasmic filaments (Fig. 2o). Average molecular velocity ranged from 1.9-2.7 μm/s and acceleration or deceleration (known as absolute acceleration) to 58-79 μm/s^2^, with no significant difference between import and export when the NPC is considered together; however, on segregation of NPC subdomains, import is more highly dynamic once transported to the nuclear basket domain and conversely export once transported to the cytoplasmic filaments (Fig. 2p,q; Extended Data Fig. 5). This is suggestive that cargo dispersal from the NPC after successful transit through the channel is faster than initial cargo interaction with NPCs prior to transport irrespective of direction.

Direction-specific tracks could further be segregated into those that underwent successful (defined as ending in the alternate compartment) or abortive transport (Extended Data Fig. 3a). The proportion of tracks that underwent successful nucleocytoplasmic transport was 29.8% for nuclear import and 36.7% for export (Extended Data Fig. 3b). As expected, abortive events were associated with lower molecular densities further into the NPC than successful events – for nuclear import this meaning lower molecular density after the cytoplasmic filaments in central and nuclear domains, and vice versa for export (Extended Data Fig. 3c,d). Interestingly, this is accompanied by trends or significance to increased dynamics of abortive trajectories during nuclear import and reduced during export (Extended Data Fig. 3e,f). Together these data demonstrate the effective application of a new technique to define single molecule nucleocytoplasmic transport dynamics at the individual nuclear pore and subdomain level in intact non-genetically modified cells. To our knowledge this is also the first determination of differential dynamics in passive nuclear import versus nuclear export, enabled through use of an intact equilibrated system.

**Fig. 3.**
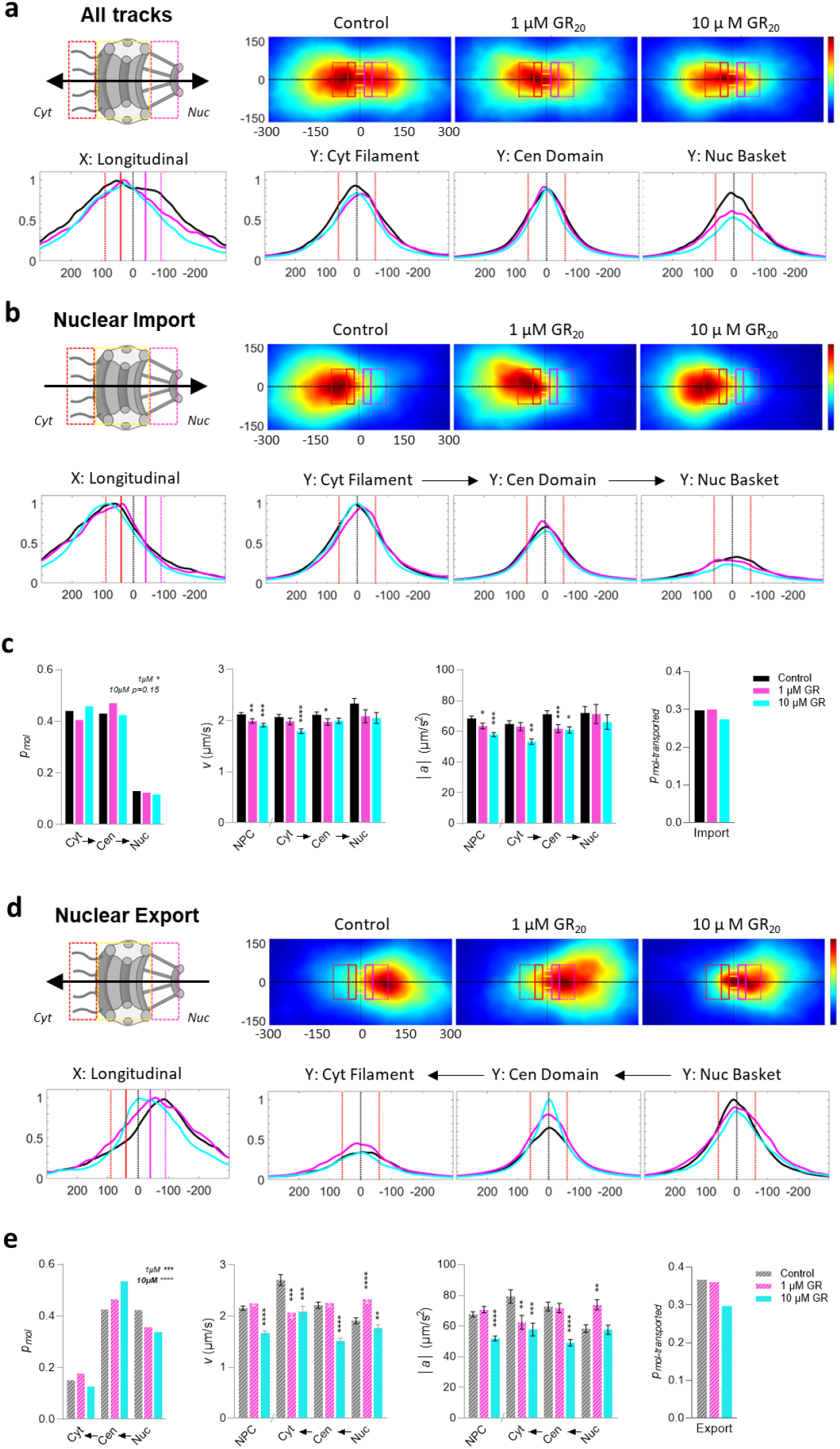
Alterations to single molecule nucleocytoplasmic transport dynamics at the nuclear pore complex subdomain level upon exposure to disease-associated toxic peptide polyGR. Neuroblastoma cell line SH-SY5Y was incubated with a small inert CF640R-tagged 10 kDa dextran cargo untreated (control - black) or with acute treatment (1 hr) of polypeptide GR20 (1 µM – magenta, or 10 µM - cyan). Each dataset is presented with a nuclear pore complex (NPC) schematic with direction of transport overlaid with subdomain dimensions with dashed lines highlighting the cytoplasmic filament (red), central (yellow) and nuclear basket (pink) subdomains. Datasets for all tracks (a) and segregation by nuclear import (b,c) and nuclear export (d,e) are presented. Each data set includes (axis in nm) density heatmaps for molecular positions within tracks with overlaid NPC dimensions; comparative longitudinal (y = 0 in respect to heatmap) density profile through the NPC; comparative transverse profiles through the midpoint of each of the cytoplasmic filament (x = −90nm), central (x = 0) and nuclear basket(x=90nm) subdomains. Quantitative analysis of nuclear import (c, solid bars) and nuclear export (e, diagonal striped bars) display data for proportional molecular density between different subdomains (pmol), velocity (v) and absolute acceleration (|a|) for total NPC or subdomains with arrows highlighting direction of transport, and proportional successfully transported tracks (pmol - transported). Statistical analyses of molecular data: proportional molecular density – chi squared vs control, velocity and absolute acceleration - one-way ANOVA with Dunnett’s multiple comparison vs control on log-transformed data for total NPC, 2-way ANOVA with Dunnett’s multiple comparison vs control on log-transformed data for subdomains; *p<0.05, **p<0.01, ***p<0.001, ****p<0.0001; N>1000 for total NPC, N>145 for subdomains; raw numerical data in Source Data File for Fig. 1.

### *Disease C9ORF72* aberrant polypeptide polyGR exerts a greater disruption on passive nucleocytoplasmic export dynamics than import

Interactome studies have demonstrated that the *C9ORF72* toxic arginine rich DPRs bind to low-complexity domain containing proteins, particularly those involved with the formation of membraneless organelles, including components of the NPC central pore domain ^24-26^. This data aligns with findings that NPC components can modify DPR toxicity in yeast, fruit fly and mammalian cell models ^44-47^. However, direct measurement of how nucleocytoplasmic transport is altered from this aspect of disease has not yet been determined. Thus, using our novel technique, we set out to test whether the *C9ORF72* DPR polyGR could alter passive nucleocytoplasmic transport dynamics. The neuron-like cell line SH-SY5Y was incubated with synthetic polyGR peptide (GR_20_) at 1 or 10 μM for 1 hour – a concentration range known to elicit toxicity under prolonged exposure ^20^, prior to tracking of the CF640R tagged 10 kDa dextran. By restricting treatment to an acute timeframe, we aimed to examine early perturbations in nucleocytoplasmic transport prior to the onset of peptide-induced cell death. In line with this, the number of tracks per cell remained unchanged (Table 1), indicating that the acute exposure conditions did not generate measurable toxicity within the period of analysis. Analysis of all tracks show that peptide treatment altered interaction with the NPC, particularly in the nuclear basket domain (Fig. 3a). Separation of directionality revealed that polyGR could impact both nuclear import and export, however, in almost all aspects export was affected to a significantly greater degree. Molecular density heatmaps display little alteration to nuclear import upon polyGR application, also reflected in the intensity line scans (Fig. 3b). However, a significant difference was detected in the molecular distribution with 1 μM GR_20_ with a 3.4% reduction in the cytoplasmic filament domain and 4.1% increase in the central domain, but this was not recapitulated or exacerbated in the higher 10 μM concentration (Fig. 3c). A similar but larger, more significant and concentration-dependent trend was observed for polyGR on nuclear export, with overt reduction of molecular density in the nuclear basket (6.6% decrease at 1 μM, 8.6% at 10 μM) and accumulation in the central channel domain (4.0% increase at 1 μM, 11.0% at 10 μM; Fig. 3d,e). These alterations are also apparent in the transverse density line plots through the respective NPC subdomains (Fig. 3b,d). Upon separation of successfully transported tracks from abortive events, trends in export dysfunction become more overt in all domains and in the nuclear basket defect during import (Extended Data Fig. 4a,b). This analysis reveals an additional difference during export, where abortive tracks have an increased molecular density in the central domain whereas successful tracks show no change in the peak molecular density but a narrower distribution, indicating molecular flow through a narrower channel upon polyGR exposure.

**Fig. 4.**
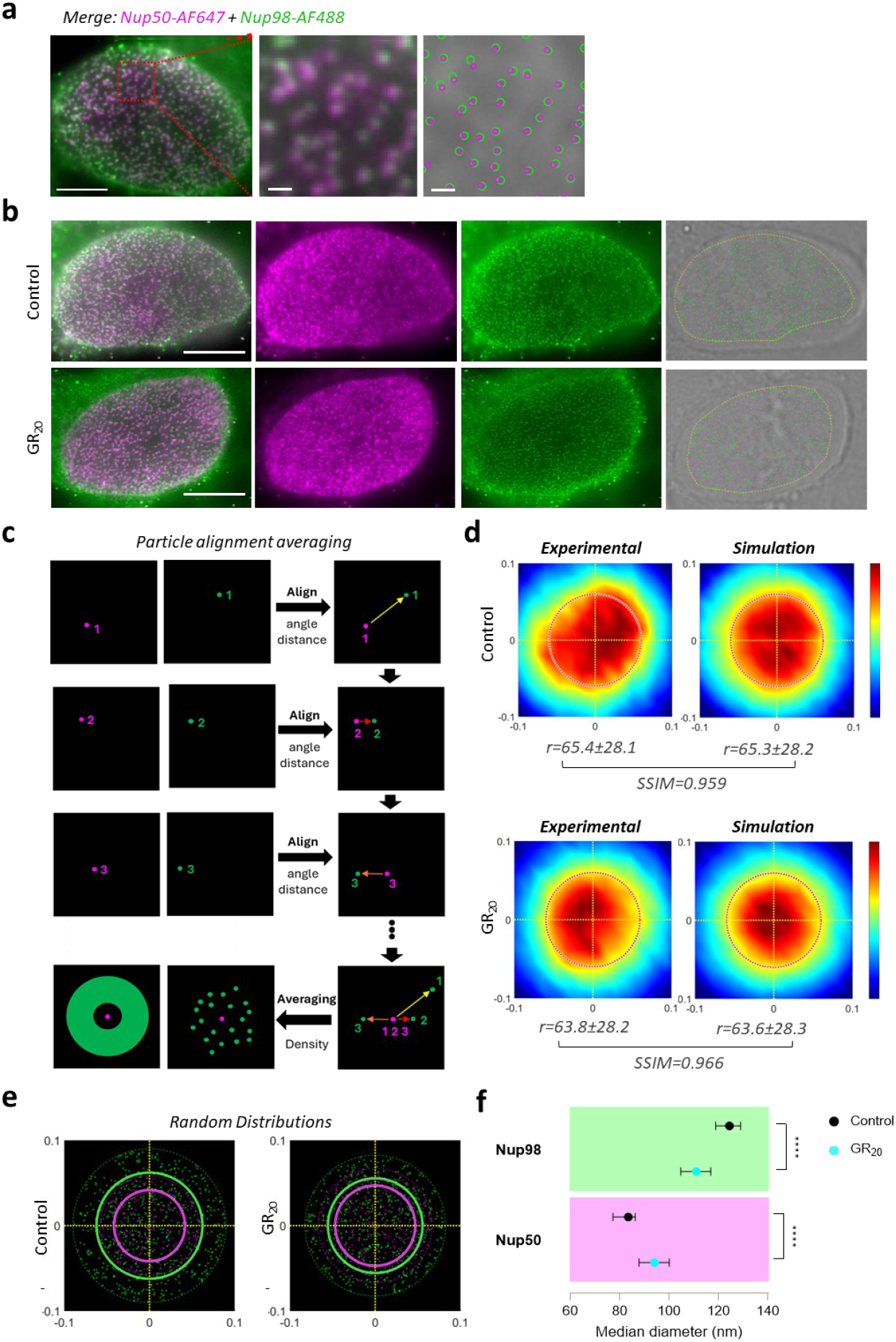
Paired localisation and diameter simulation analysis for NPC structure. Neuroblastoma cell line SH-SY5Y untreated (control) or with acute treatment (1hr) of polypeptide GR20 (10µM), as per single molecule tracking experiments, and then immunostained with dye-conjugated antibodies for the NPC components Nup98 and Nup50. a) Example fluorescence image of improved immunostaining detection of individual NPC spot detection of AF647-labelled Nup50 (magenta) and AF488-labelled Nup98 (green) acquired using HILO microscopy, with enlarged view and respective definition of Nup50/98 positions; scale bars are 5 μm and 500nm for enlarged views. b) Representative fluorescence images of untreated (control) and polyGR-treated cells with output spot detection for Nup98 and Nup50 within deep learning nuclear envelope central boundary definition (yellow dashed line); scale bars are 5 μm. c) Schematics of process for paired particle localisation and alignment averaging for Nup98 positioning (green) in reference to Nup50 (magenta). d) Output density heatmap of Nup98 positioning in reference to Nup50 from paired localisation analysis for experimental data of control and polyGR-treated cells with comparative best fit simulation heatmap, with values for mean radii – here, indicating the distance between Nup50 and Nup98 - (±SD) and structural fidelity between experimental and best fit simulation data using the structural similarity index (SSIM) below. e) Best fit simulated distribution of Nup50 (magenta) and Nup98 (green) – now reflecting their effective position in a nuclear pore complex - used to generate heat map in (d) with lines for respective median Nup98 and Nup50 diameter in control and polyGR treated conditions. (f) Quantification for determined median diameter (±95% CI) of Nup98 and Nup50 under control and polyGR-treated conditions using paired localisation and diameter simulation analysis. Statistical testing: unpaired Mann Whitney; ****p<0.0001; N>250; raw numerical data in Source Data File for Fig. 3.

Assessment of subdomain-specific NPC transport dynamics demonstrate poly-GR induced alterations to both molecular velocity and absolute acceleration (acceleration/deceleration), with trends or reaching significance to ∼10% average decrease overall and in most subdomains during both import and export, but again more overt during export (Fig. 3c,e; Extended Data Fig. 5). Notably, the success rate of the measured passive nucleocytoplasmic transport events through NPCs reflects the altered dynamics with nuclear import showing a reduction of 8.1% upon 10 μM polyGR, and export a reduction of 19.1%. Interestingly, when single molecule localisations are assessed within the whole nucleus (up to central boundary of nuclear envelope), polyGR leads to an increase at both 1 and 10 μM (Extended Data Fig. 6a,b). Thus, although passive nuclear export dynamics and transport success rate are reduced significantly more than import, the 10 kDa dextran cargo undergoes nuclear accumulation with polyGR, in agreement with previous studies in permeabilised cells ^48^. This is likely due to polyGR-induced retention of cargo through association with nuclear structures upon nuclear import, such as the nucleolus ^20, 49-51^. Indeed, the decline in nuclear molecular counts during the experimental acquisition in polyGR treated conditions is in agreement with nuclear molecules being more static which renders them susceptible to photobleaching, rather than being dynamically displaced and replaced with new molecules within the plane in untreated cells (Extended Data Fig. 6c).

Together, through our single molecule tracking methodology we have been able to show that soluble disease-associated *C9ORF72* polyGR peptides can disrupt the molecular flow of inert molecules through NPCs. Upon peptide exposure, inert cargo interacts less with nuclear basket and cytoplasmic filament structures and more with the central channel, all with reduced dynamics of interactions. This results in a reduction in bidirectional nucleocytoplasmic transport of this cargo, with more overt changes in nuclear export. However, summative analysis shows nuclear accumulation, potentially via association with nuclear structures, again with reduced molecular dynamics. These differences highlight the necessity to study nucleocytoplasmic transport dynamics directly at the single molecule resolution to overcome other factors that influence summative data.

### Paired localisation and diameter simulation analysis for NPC structure and impact of *C9ORF72* DPR polyGR

Our findings on nucleocytoplasmic transport molecular dynamics demonstrate an enriched inhibitory effect of polyGR on passive export and an associated increased interaction and narrower passage through the central pore domain and a reduced interaction with the nuclear basket domain. Thus, we next aimed to investigate whether these changes were due to an alteration in the structural organisation of these NPC subdomains. Here, we used another novel approach combining localisation analysis and simulation modelling to determine the diameter of central domain and nuclear basket components within the NPC. Immunostaining for the nucleoporins Nup98 and Nup50 were selected for representation of the central and nuclear basket NPC domains, respectively, while also having antibodies with the specificity required for single molecule localisation microscopy. Cells underwent the same cellular treatments prior to fixation and immunostaining as single molecule tracking experiments, after which cells were pressed to flatten the nuclear envelope closer to the coverslip for improved NPC image acquisition (Fig. 4a,b and Extended Data Fig. 7a). We set the maximum pairing distance for these nucleoporins at 120 nm, equivalent to the outer diameter of the human nuclear pore, to prevent erroneously grouping of nucleoporins from separate NPCs; ∼80% of single molecule localizations fell within this distance (Extended Data Fig. 7b,c).

In control cells, the densities of Nup98 and Nup50 were 4.1 and 3.1 spots per μm^2^, respectively, with a 77.8% pairing frequency and an average inter-spot distance of 68.0 nm (Extended Data Fig. 7d-f). Acute polyGR peptide treatment (10 μM for 1hr) resulted in no significant difference in the Nup98 or Nup50 density at the nuclear envelope and pairing frequency was maintained at 77.3%; however, average inter-spot distance showed a slightly reduced trend (down to 64.5 nm), suggesting potential nuclear pore structural rearrangement (Extended Data Fig. 7f). To give further definition of these changes, a combination of single molecule localisation alignment and a data simulation approach was used to predict the diameter of the nucleoporins within NPCs from their relative positions. Firstly, the single molecule localisations were aligned in both angle and distance, represented in a density heatmap of Nup98 positioning in comparison to Nup50 (Fig. 3c,d). Then, random spot distributions were generated to simulate Nup50 and Nup98 positions, with optimal distribution selected based on fit with experimental inter-spot measurements; this achieved a high structural fidelity between the modelled and experimentally measured configurations of nucleoporins within NPCs (structural similarity index, SSIM ^52^ >0.95) (Fig. 4d,e). Using this approach, we could determine that in control cells Nup98 and Nup50 are predicted to have average diameter positions of 95 and 60 nm, respectively (Fig. 4e,f). These measurements represent epitope-defined positions from relative positions derived from the fluorophore-conjugated primary antibody localisations combined with simulation and should therefore be interpreted as effective rather than absolute positions. In polyGR treated cells, the diameter of Nup98 contracted to 86 nm while that of Nup50 expanded to 71 nm (Fig. 4e,f). Importantly, these structural data align with molecular flow results upon polyGR treatment: contraction of the central domain component Nup98 correlates with the narrower passage through the central domain, and expansion of the nuclear basket component Nup50 with a reduced impediment to transport and thus reduced interaction and dwell time through this domain.

### Convergence of structure-function alterations at the NPC to disease associated TDP-43 proteinopathy

According to the results of our study thus far, the *C9ORF72* DPR polyGR can significantly disrupt both nucleocytoplasmic transport and NPC structural organisation. As already introduced, a convergent feature in ALS and FTD, also associated with the *C9ORF72* mutation, is TDP-43 proteinopathy. This nucleocytoplasmic shuttling RNA-binding protein becomes mislocalised from a predominantly nuclear localisation to the cytoplasm in disease where it accumulates and forms a classical inclusion pathology ^11^. Interestingly, TDP-43 has been proposed to undergo receptor-mediated nuclear import but receptor independent, passive export ^53-55^. Thus, we investigated whether altered TDP-43 cellular distribution is affected in our model of early *C9ORF72* DPR NPC pathology. Again, under the same conditions as nucleocytoplasmic transport dynamics and NPC structure analyses (10 µM polyGR, 1 hr), cells were immunostained for TDP-43 and imaged using a tilted beam angle (7° below the incident) to provide more accurate signal to noise. While no overt cytoplasmic mislocalisation was visible from acquired images, quantitative summative analysis revealed a significant 16.0% reduction in the nuclear to cytoplasmic ratio upon polyGR treatment, that appears to be driven predominantly by a trend in reduced nuclear protein level (12.1%, p=0.17) with a subtle non-significant increase in the cytoplasm (5.5%, p=0.35), which is also apparent from frequency distributions of intensity values (Fig. 5a-c). This indicates that acute disruption of nucleocytoplasmic transport upon polyGR exposure includes early mislocalisation of TDP-43.

**Figure 5.**
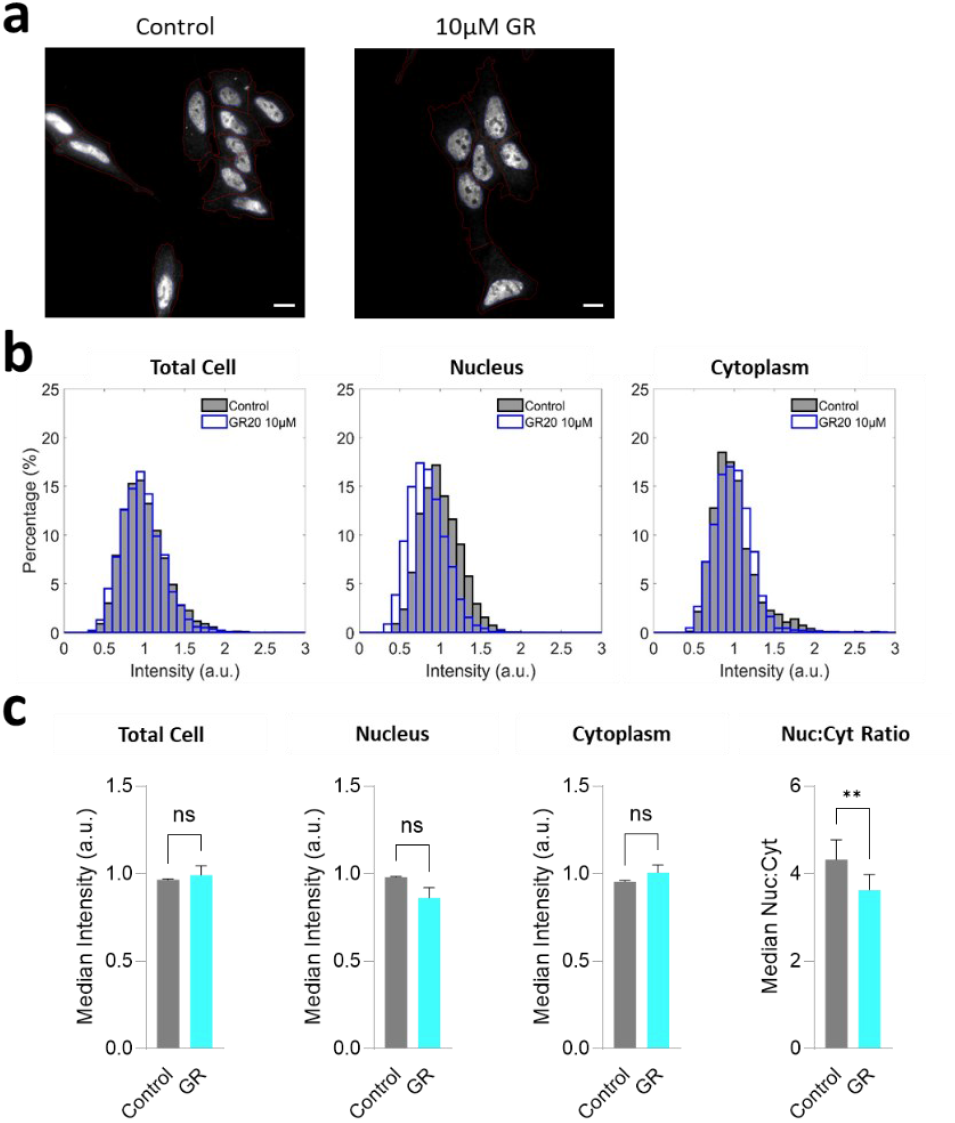
Early indication of TDP-43 cytoplasmic redistribution in cells acutely exposed to disease-associated peptide polyGR. Cytoplasmic mislocalisation of the RNA-binding protein TDP-43 is a key hallmark in C9ORF72-associated, and other, forms of sporadic and familial ALS/FTD. Here, TDP-43 level and cellular distribution was assessed through immunostaining of TDP-43 of SH-SY5Y cells following treatment with GR_20_ peptide (10µM, 1hr), as per single molecule tracking and NPC structure analysis experiments. a) Representative fluorescence images of control and polyGR-treated cells; scale bars are 10 µm. b) Histograms of intensity values normalised to the mean of the control per biological replicate, and (c) median values for normalised total cell, nucleus and cytoplasm compartments, and nuclear-to-cytoplasm ratio (non-normalised values). Statistical testing: paired or ratio paired (for Nuc:Cyt) t-test; N=3; **P<0.01; raw numerical data in Source Data File for Data Figure 5.

Together, the integration of new methodologies for single-molecule tracking of passive nucleocytoplasmic transport in intact, unmodified cells, with structural analyses based on adapted conventional immunostaining, paired particle localisation and simulations delineates previously unappreciated features of nuclear pore functionality. We apply these to unveil disturbances to nuclear pore function and architecture from exposure to the *C9ORF72*-associated aberrant disease peptide polyGR, with a preferential vulnerability of the nuclear basket and central channel and correspondingly greater effect on passive nuclear export than import. These defects coincide with the emergence of TDP-43 proteinopathy, establishing a mechanistic link between polyGR-mediated nuclear pore perturbation and disease-relevant cellular pathology (Figure 6).

**Figure 6.**
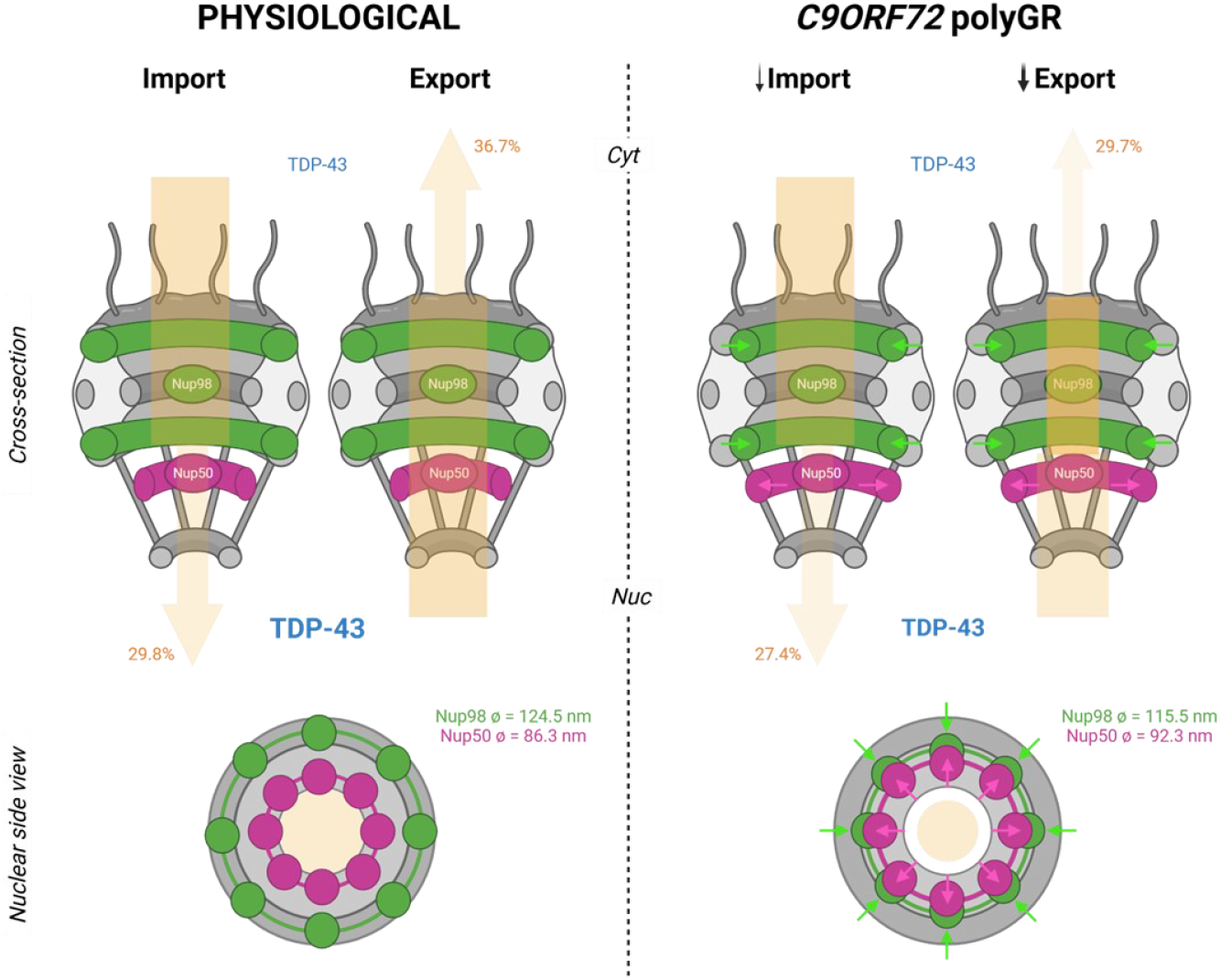
Schematic summarising disease-related findings unveiled by novel techniques for single molecule nucleocytoplasmic transport tracking and nuclear pore structural organisation. Nucleocytoplasmic transport (beige) in physiological and C9ORF72 polyGR (10 µM) disease conditions with width and density of arrow denote transport passage and molecular density, and percentage success rate of tracks. Nuclear pore structure schematic in grey in cross section (upper) and nuclear side view (lower) with Nup98 positions denoted in green and Nup50 in magenta, arrows highlighting change in organisation upon exposure to polyGR, and values for simulated diameter measurement. Associated change in nucleocytoplasmic distribution of disease-associated protein TDP-43 shown in blue. Created in BioRender.

## Discussion

Here, we have presented two novel techniques to resolve the structure-function relationship of the large multiprotein NPC in intact live cells, and their application to understand neurodegenerative disease biology. Firstly, we developed a method for single molecule nucleocytoplasmic transport tracking in intact and non-genetically modified cells with resolution of NPC subdomain-specific dynamics. Secondly, we brought together standard immunostaining with paired single molecule localisation and simulation analysis to determine the ring-based structural organisation of components of the NPC, without the requirement for specialised super-resolution microscopy. We also used higher resolution imaging to detect the very early stages of a hallmark pathology associated with NPC structure-function in disease. Together, these methodologies have revealed that the *C9ORF72* DPR polyGR peptide can disrupt passive nucleocytoplasmic transport, particular export, with altered structural organisation and molecular interaction particularly within the central pore and nuclear basket domains of the NPC, and that these alterations can initiate disruption of TDP-43 nucleocytoplasmic homeostasis.

We introduced several novel features into our single molecule nucleocytoplasmic transport tracking approach – the ability to use live non-genetically modified intact cells, machine learning to determine the nuclear envelope position from brightfield images, and the prediction of NPC localisation to map interactions. This use of non-genetically modified cells, both for NPC and nuclear envelope localisation, prevents any modification of structure and dynamics by fluorescent protein labelling. We also prioritised the use of intact live cells to study nucleocytoplasmic transport dynamics in a fully physiological environment. One of the key challenges in the use of live cells was the ability to image at depth in a biological sample in order to acquire data at the nuclear envelope (∼2 μm in most cell lines including SH-SY5Y). This is been traditionally approached by studying isolated nuclei ^33-35^, however, removal of the cytoplasm and factors that undergo transient interaction with NPCs is known to impact nuclear pore integrity and dynamics ^36, 37, 56^. We overcame this by optimising parameters for reducing and filtering noise, including fluorescence bleaching at the level of the coverslip and post-acquisition noise filtering, combined with use of sensitive detectors for low signal range. More recently alternative imaging approaches have been applied to assess the molecular dynamics of nucleocytoplasmic transport in intact cells, including single-point edge-excitation sub-diffraction (SPEED) microscopy ^57^ and MINFLUX ^58^. SPEED microscopy enables millisecond tracking of single molecules passing through individual NPCs in live cells, but measurement is at the single NPC level. MINFLUX achieves nanometre localisation precision for single molecule tracking but has demanding optical requirements and also a small field of view. In contrast, our approach enables simultaneous single molecule tracking across the entire nuclear envelope in intact live cells, providing a global readout of NPC permeability and dysfunction, while still enabling single nuclear pore subdomain resolution. These distinctions underscore how different imaging modalities capture complementary scales of nucleocytoplasmic transport. Our method therefore occupies a unique niche by enabling and higher throughput live cell, nucleus-wide transport profiling without genetic modification or specialised optical equipment.

Using our single molecule nucleocytoplasmic transport tracking methodology, we could determine attributes of nucleocytoplasmic transport of an inert cargo (passive, receptor-independent transport) including molecular dwell time, velocities and acceleration through individual NPC sub-domains. To the best of our knowledge no studies have determined the kinetics of passive transport through the NPC, however, studies of receptor-mediated (termed “active”) and passive transport in permeabilised cells have demonstrated interaction/transport through NPCs generally in the range of a few milliseconds ^59-65^ but sometimes up to seconds ^63, 66^. Similarly, more recent studies of actively transported cargo also measure transport in the range of milliseconds ^57, 58^. Here, we detected molecular movement of a 10 kDa dextran through NPCs at an average of ∼2 μm/sec, which, taking into account a 180 nm transversal length of an NPC, equates to an NPC interaction time of ∼90 milliseconds. This is lower than previous reports for a 10 kDa dextran in permeabilised cells ^60, 61^, which may reflect slower passive transport impeded by significant transiently interacting factors in an intact cell system, as shown for increasing transport receptor occupancy ^61^.

We could also determine that the successful passage of our 10 kDa inert cargo was higher for basal nuclear export (36.7%) than import (29.8%), which would not be expected of a fully passive transport mechanism. However, no molecule is fully inert, and the NPC is an asymmetrical structure across the nuclear envelope, thus this directional difference may reflect a higher cargo-NPC interaction rate with the long cytoplasmic filaments of NPCs but without conversion to complete NPC translocation compared to a more limited and selective cargo interaction with the nuclear basket. Reported success rates for actively transported cargo vary widely (∼20-68%) ^57-59, 62-66^ which may reflect the different cargos or experimental approaches. A previous study using 10 kDa dextran measured 51 % success rate for both passive import and export ^60^, again this difference may reflect the absence of intrinsic factors in permeabilised cells, or experimental approach. However, our rates are within the range of receptor-mediated as passive transport, in line with the proposed continuum of passive and active transport dependent on size and surface properties of cargo ^4^. A limitation of our success rate measurements is that due to the prediction of NPC localisation, non-transported tracks could include molecules that exhibit NPC-interaction behaviour but actually are only interacting with the membrane. This is mitigated by the stringent NPC-interaction criteria applied and thus is unlikely to account for a significant proportion of tracks.

In order to correlate functional changes at the NPC with structure, we also developed a method for resolution of the ring-based organisation of endogenous NPC components using optimised conventional immunostaining, dye-labelled primary antibodies and angled-beam imaging. This does not offer the resolution of higher super-resolution methodologies, but these generally require more specialised reagents and are significantly more time consuming. Using this approach, we could define the paired distance between the central domain component Nup98 and nuclear basket component Nup50. Then, using simulations for comparative location and fitting with experimentally defined parameters, we could define Nup98 and Nup50 diameters at NPCs in untreated cells as 95 and 60 nm, respectively. The exact radial positions of Nup98 and Nup50 in intact cells are not fully defined. However, Nup98 is known to anchor to the inner ring scaffold at both nuclear and cytoplasmic faces of the NPC at a diameter of ∼60 nm ^67^, but with projections of its unstructured region into the central channel and also potential into nuclear basket and cytoplasmic filament domains meaning its position could range from 0->120 nm ^68^, and is also a moderately dynamic nucleoporin ^69^. Nup50 is localised to the nuclear ring anchor of the nuclear basket which has a diameter of ∼80 nm, but is a highly dynamic nucleoporin ^69^ and thus likely to adopt positions within the entire nuclear basket structure, diameter of ∼40-100nm. Thus, the accuracy of our measurements is difficult to determine, but they are both within the predicted range. The limitation of application of this technique for the NPC will be determined by the availability of antibodies that provide enough specificity for single molecule localisation microscopy.

These approaches were applied to gain pathomechanistic insight into the neurodegenerative diseases ALS/FTD. Nucleocytoplasmic transport has been extensively implicated in these disorders and particularly linked to toxicity associated with DPR peptides produced from the most common mutation in *C9ORF72*. However, studies in this field have been limited to indirect measurement of transport with assessment of change in nuclear and cytoplasmic levels, mostly statically, but occasionally with time-course. Alterations to the composition of NPCs have been detected in *C9ORF72*-disease, but mainly in association with DPR-independent mechanisms ^70, 71^. In our studies, we did not detect alterations to NPC number and intensity data suggested no loss of Nup98 or Nup50. However, our model uses only acute DPR application, which may represent an early stage in disease prior to loss of NPC components or an effect that occurs in parallel in the disease context. We found that the *C9ORF72* DPRs could alter both passive nucleocytoplasmic transport dynamics and NPC structural organisation. Indeed, structural alterations aligned well with functional changes. An enlargement in the diameter of the nuclear basket component Nup50 could reduce the probability of molecular interactions in this domain and thus hinder transport less resulting in the reduced molecular dwell time observed during nuclear export. The DPR polyGR also caused constriction of the central pore component Nup98 which could explain the passage of molecular flow through a narrower channel. It is currently unknown whether these effects occur through direct binding of the arginine rich DPRs – supported by interactome data^24-26^ and one study reporting polyPR localisation to NPCs in isolated nuclear envelopes of giant nuclei from *Xenopus laevis*^72^. However, changes could also occur via indirect effects either from different NPC domains or protein interactions or non-NPC cellular interactions. Recent studies have demonstrated that arginine rich DPRs can directly bind to nuclear import receptors reducing their availability for receptor-mediated transport ^28, 73^, demonstrating a mechanistic link between DPRs and nucleocytoplasmic transport disruption, but one independent to the receptor-independent transport disruption described here.

Finally, we determined that the polyGR-induced disruption of NPC structure-function we observed using our novel techniques, was also correlated with early features of TDP-43 proteinopathy. The *C9ORF72* mutation and expression of polyGR has been identified to cause TDP-43 proteinopathy in cell and animal models ^29, 74^. Thus, our findings corroborate that polyGR peptides can initiate this step even in acute settings. Although the here reported changes in passive nucleocytoplasmic transport dynamics may contribute to the altered TDP-43 nucleocytoplasmic homeostasis observed, its direct contribution cannot yet be determined. Indeed, its nuclear loss would be expected to be associated with increased propensity for passive nuclear export rather than reduced, and its nuclear import is predominantly control via receptor-mediated transport ^75^ where impairment of passive import would not be predicted to impart a significant effect. Interestingly, although single molecule transport dynamics were reduced for both import and export by polyGR, summative analysis revealed nuclear accumulation, which could be explained by the greater effect on export than import, but may also occur due to binding to nuclear structures, such as the nucleolus, which the arginine rich DPRs are also known to affect ^20, 49, 51^. Thus, again these disparities and contributions of other processes highlight the importance of studying nucleocytoplasmic transport directly using single molecule methodologies, rather than relying on inference from summative assessments.

In conclusion, we here present two novel methodologies for the determination of NPC functionality in live, unmodified cells and NPC structure using conventional immunostaining and angled imaging, which can be used to gain novel insight into physiological nucleocytoplasmic transport dynamics and how this becomes dysfunctional in disease.

## Methods

### Reagents

Aberrant *C9ORF72* polyGR peptide – GR_20_ – was synthesised by Cambridge Research Biochemicals with > 95% HPLC purity. Unlabelled polyGR was used to avoid any interference from dye-associated electric charges. mCherry-LaminB1-10 was a gift from Michael Davidson (Addgene plasmid #55069).

### Cell Culture and Treatment

SH-SH5Y cells were maintained in a humidified incubator at 37 °C, with 5% CO_2_ in cell medium: DMEM-F12 (gibco) with 15 % FBS, 1 % GlutaMAX, 1 % Penstrep, 1 % Sodium pyruvate and 1 % non-essential amino acids in a controlled humidified, 5% CO_2_, 37 °C incubator. Cells were plated onto µ-Slide 8-well high 1.5H glass bottomed chamber slides (ibidi) without coating. For single molecule tracking, the cells grew on chamber slides for 24 hours prior to 1 hour treatment with polyGR peptide, if required. For single molecule tracking experiments, 1 mg/mL CF640R-tagged 10 kDa dextran solution (Cambridge Bioscience) was added to cell media by gently applying the solution to the side wall of the well (final concentration - 50 µg/mL), maintained for several minutes, and washed by replacing with fresh growth medium. Immediately before the experiment, the medium was changed to transport buffer (20mM HEPES, 110mM KOAc, 5mM NaOAc, 2mM MgOAc, 2mM DTT, pH 7.3 ^66^).

### Transfection

For deep learning training, cells were transfected with mCherry–LaminB1-10 plasmid by electroporation. Cells were dissociated using trypsin and neutralised with complete growth medium. Following cell counting, approximately 1 × 10^5^ cells per condition were collected by centrifugation at 1200 rpm for 4 min, washed once with PBS (1200 rpm, 5 min), and resuspended in nucleofection buffer. For each reaction, cells were resuspended in 82 µL nucleofection buffer supplemented with 18 µL electroporation enhancer buffer (Invitrogen, MPK10025), and 1–1.3 µg plasmid DNA was added immediately prior to electroporation. Electroporation was performed following standard procedures for mammalian cells. Following electroporation, cells were immediately transferred into pre-warmed complete growth medium and allowed to recover under standard culture conditions prior to downstream imaging.

### Single Molecule Transport Image Acquisition

We used a Nikon Eclipse TiE inverted fluorescence microscope equipped with high-NA oil immersion objectives (Nikon CFI Apo TIRF 100XC Oil NA1.49 and Nikon CFI SR HP Apo TIRF 100XAC Oil NA1.49), TIRF illuminator, multiple lasers and cameras (Andor iXon EMCCD and ORCA-Flash4.0 Digital CMOS).

For single molecule imaging, we first reduced noise using TIRF (total internal reflection fluorescence) to photo bleach molecules close to the coverslip (67.55° using the 647nm laser, 1 min). We then used a HILO (Highly Inclined and Laminated Optical sheet) imaging approach, which uses a low-powered laser at below-critical angle to increase illumination of sample further above the coverslip while maintaining selectivity of single molecules. This was optimised to 1.67° below the critical angle, giving a final angle of 59.56°, for acquisition at a depth of approximately 2 µm above the coverslip and at the lower edge of the nuclear envelope, with some adjustment for focus by further reducing the incident laser beam angle, at a rate of 53 fps and acquisition of 10,000 frames (∼3 minutes).

### Data Analysis Code

Code for analyses protocols can be found at the Github of Dr Seoungjun Lee and the Mizielinska Lab (www.github.com/MizielinskaLab).

### Single Molecule Spot detection and Molecular Tracking

Single molecule images were denoised to minimize noise from the EMCCD camera and increase the contrast of molecules using CANDLE (Collaborative Approach for Enhanced Denoising under Low-light Excitation) in MatLab ^41^. Spot detection and molecule tracking were then analysed using TrackMate ^76^, custom Python and MatLab code. We used spot detection for LoG (Laplacian of Gaussian particles) detection based on local maxima and suitable for Gaussian-like spots under noisy conditions ^77, 78^. The detected spots were examined for quality with the local maximum value, and spots were immediately removed if the detection quality was lower than the detector value. Since the detected spots for transport through NPCs is fast and very variable, which makes them unsuitable for LAP trackers, we applied the nearest neighbour search tracker ^79^. For quality control of tracking, we excluded molecular clusters, out-of-focus molecules, molecules with overlapping tracking locations (<160 nm), and molecules with overlapping tracking times (<1 s).

### Deep learning for prediction of the nuclear envelope position

All deep-learning analyses were performed on a Linux workstation equipped with 512 GB of system RAM, four NVIDIA Titan RTX GPUs, and an Intel Xeon CPU. The training dataset consisted of paired images comprising mCherry–LaminB1 fluorescence images and the corresponding brightfield images. Increasing the number of input channels and patch sizes improved agreement between the trained model and the predicted images, however, this came at the cost of slower processing and substantially increased demands on GPU memory and system RAM. For model training, we employed a 2D FNet architecture with 256 multi-channels, 4 depths, 512×512 patch sizes, 4 batch sizes, 1,000 buffer sizes, and 18,000 buffer switch frequencies. These parameters were selected to balance model performance and computational feasibility.

Our deep-learning framework was adapted from a previously described label-free fluorescence prediction approach^42^ and optimized to predict the position of the nuclear envelope (NE) from brightfield images. The model was trained using paired mCherry-LaminB1 fluorescence images and brightfield images, where the brightfield image served as the input and the corresponding fluorescence image as the target output. The trained model generated fluorescence-like predictions from brightfield inputs that closely matched the ring-shaped NE signal in the ground-truth mCherry-LaminB1 images (Extended Data Fig. 2). Compared to a previously established method^66^, our deep-learning approach provides enhanced spatial accuracy as it resolves the central NE position as well as regions above and below the focal plane. This is particularly important when imaging near the bottom of the cell, where traditional analysis methods frequently misassign NE position. For the training, each image was contributed to by 1,000 brightfield frames. These frames were merged into a single high-contrast composite image to enhance the visibility of the NE, which is not readily detectable in individual brightfield frames. Once the lamin B1 positions were defined, a 29 nm outward adjustment was applied for the definition of the nuclear envelope central position, accounting for lamin B1 localisation within the lamina at the inner nuclear membrane interface. This was approximated based on the difference between the reported midpoint of inner–outer nuclear membrane spacing (40-50 nm^80^) and the midpoint of the nuclear lamina meshwork (10-15 nm^81^), of which Lamin B1 is a component. During SMT acquisition, we applied the same brightfield imaging and processing pipeline. For each image sequence, 1,000 brightfield images were acquired before and another 1,000 after SMT imaging. Each set of 1,000 frames was processed into a composite brightfield image, which was then used as input for NE prediction by the deep-learning model. Because minor cell drift can occur during SMT acquisition, the NE position for each cell was calculated as the midpoint between the NE positions predicted from the before SMT and after SMT composite images. If the displacement between these two predicted NE positions exceeded 600 nm, the cell was excluded from further analysis to ensure that only cells with minimal movement were retained for accuracy of NE localization and SMT mapping.

### Trajectory alignment to the nuclear envelope and NPCs

Single molecule tracks were overlaid on the defined nuclear envelope central position. Single molecule tracks that did not intersect with the central location of the nuclear envelope, as determined by deep learning were excluded. As the nuclear envelope provides a significant barrier to nucleocytoplasmic transport, transversal across the NPC will be the principal route of transport of our small inert cargo. Thus, after rotation in line with the nuclear envelope, single molecule tracks were overlaid on an NPC domain model of known mammalian NPC dimensions ^82, 83^ (Fig. 1). We stringently defined track interaction with the NPC (and not low affinity interactions with the nuclear envelope), only including tracks with ≥10 molecular positions aligned with NPC dimensions and ≥3 positions with the NPC central pore domain.

Data is presented as all tracks and all NPC interaction or separated into: direction - import and export – defined by starting points; subdomains – nuclear basket, central domain, and cytoplasmic filaments (Fig. 1); successful or abortive – defined by end point beyond alternative compartment to the starting point (Extended Data Fig. 3). Molecular heat maps, normalised line scans and quantitative data for molecular dwell, velocity and absolute acceleration were generated in MatLab. Success rates were calculated by the proportion of tracks that were successfully transported (beyond the nuclear basket for import, and the cytoplasmic filament domain for export) in comparison to total track count for the associated direction.

Data was also collected for all tracks that passed initial quality control for cumulative single molecule nuclear accumulation and dynamics (within defined NE, not interacting with NPC).

### Immunostaining

For nuclear pore structure analysis immunostaining using dye-conjugated antibodies was performed. SH-SY5Y cell in 8-well chamber slides were once washed with PBS and rinsed with 200 µl 4% PFA (ThermoFisher Scientific, #043368.9M) in PBS. The cells were permeabilized with 200 µl 0.4% Triton X-100 in PBS for 3 minutes and fixed with 200 µl 4% PFA in PBS for 15 minutes. To improved immunostaining, cells were pressed with a plastic coverslip using a tweezer then removed (Extended Data Fig. 6a). Cells were washed three times with PBS and blocked with 5% BSA in PBS for 1 hour at room temperature, then incubated overnight at 4°C with conjugated antibodies (1:50) freshly diluted in blocking buffer, including anti-Nup50 (G-4) Alexa Fluor 647–conjugated antibody (sc-398993 AF647, Santa Cruz Biotechnology) and anti-Nup98 (C-5) Alexa Fluor 488–conjugated antibody (sc-74578 AF488, Santa Cruz Biotechnology). After overnight incubation, cells were rinsed three times in PBS for 5 minutes and fixed again with 4% PFA in PBS for 15 minutes, washed three times with PBS and stored in PBS at 4 °C prior to image acquisition.

### Pairing assays image acquisition and analysis

We investigated the pairing between Nup98 and Nup50 using immunostaining of SH-SY5Y cells with or without treatment with 10 μM GR_20_ peptide for 1 h, as before. Prior to imaging, the medium was changed to OxEA buffer (PBS, 50 mM β-Mercaptoethylamine hydrochloride, 3% OxyFlour, 20% sodium DL-lactate solution, pH 8–8.5) to enhance molecular blinking and brightness^84^. AF647 labelled Nup98 and AF488 labelled Nup50 were imaged using a 647 nm laser and a 488 nm laser, respectively, on a Nikon Ti2 STORM TIRF microscope equipped with an autofocus system and a SR HP Apo TIRF 100× oil-immersion objective. To minimise the photobleaching and Interference effect on a coverslip, we used 50 ms imaging acquisition time and an 8° inclined laser beam. The exposure time and laser power of the microscope were optimized for spot resolution in Nup98 and Nup50, likely meaning that not all Nup molecules per NPC were excited, preventing saturation to improve single-molecule resolution.

Prior to single-molecule localisation analysis, denoising was performed using the CANDLE algorithm, as before. Single-molecule spots were detected using TrackMate ^76^ with a Laplacian of Gaussian (LoG) filter, a spot diameter of 195 nm (3 camera pixels), and channel-specific detection thresholds (far-red, 80; green, 100). Channel alignment between far-red and green signals was performed by defining the centre of the highest localisation density. Detection regions were manually defined based on the far-red Nup50 signal, and the selected area was reduced to 90% of its original size to minimise edge effects. The maximum pairing distance between Nup50 and Nup98 localisations was set to 120 nm, corresponding to the approximate diameter of the nuclear pore complex; localisation pairs exceeding >120 nm was excluded as they were unlikely to originate from the same NPC. Quantitative measurements included pairing efficiency, Nup spot density per area, and the inter-protein distance between Nup50 and Nup98 within individual NPCs.

### Simulation of particle alignment averaging

To give further definition of the positioning of Nup98 and Nup50 within individual NPCs, a simulation approach was used to predict the diameter of the nucleoporins within NPCs from their relative positions. Firstly, single molecule localisations of Nup98 and Nup50 were aligned in both angle and distance. Then, ∼100,000 random spot distributions were generated per simulation to model Nup50 and Nup98 positions, and ∼120,000 simulations were performed to identify the best-fit positions matching the experimental inter-spot measurements. Similarity was calculated by Structural Similarity Index (SSIM) ^52^; simulations with the closest results were presented. Simulated positioning data was then used to calculate the median diameter for control and polyGR treated analyses.

### TDP-43 analysis

SH-SY5Y cells were prepared with or without treatment with 10 μM GR_20_ peptide for 1 h, as before. Immunostaining was performed using the same fixation and permeabilisation methods as described for nuclear pore structure analysis. Cells were then washed three times with PBS and blocked with 5% BSA in PBS for 1 h at room temperature, followed by incubation with a primary antibody against TDP-43 (Proteintech, 12892-1-AP) diluted 1:500 in blocking buffer for 1 h at room temperature. After washing with PBS, cells were incubated with an Alexa Fluor 488–conjugated secondary antibody (Invitrogen, A32731TR) diluted 1:500 in blocking buffer for 1 h at room temperature, washed three times with PBS, and mounted using Fluoro-Gel mounting medium with Tris buffer (Electron Microscopy Sciences, 50-247-04). Imaging was performed using a 7° tilted laser beam, acquiring 18 × 18 large image stacks with an exposure time of 50 ms. Cell segmentation and fluorescence intensity quantification were performed using CellProfiler ^85^, and quantitative data analysis was carried out in Matlab.

## Declarations

### Ethics approval and consent to participate

N/A

### Consent for publication

N/A

### Data Availability

All data generated and analysed during this study are included in this article and its supplementary information files, or available from the corresponding authors on reasonable request.

### Authors’ contributions

Conceptualisation of study, methodology and study design, formal analysis, project administration, reviewing of draft: SL, SM. Data acquisition, writing of original draft: SL. Funding, supervision: SM.

## Acknowledgements

We thank Dr George Chennell and the team at the Wohl Cellular Imaging Centre at King’s College London for the maintenance of the microscopy facility.

## List of abbreviations

ALS: amyotrophic lateral sclerosis
FTD: frontotemporal dementia
DPR: dipeptide repeat
polyGR: polypeptide of glycine-arginine
NPC: nuclear pore complex
Nup: nucleoporin
TIRF: total internal reflection fluorescence imaging
HILO: highly inclined and laminated optical sheet imaging
CANDLE: Collaborative Approach for Enhanced Denoising under Low-light Excitation
SSIM: Structural Similarity Index

