## Supplementary Figures for "Novel single molecule imaging approaches reveal structure-function alterations to the nuclear pore complex in early *C9ORF72*-associated TDP-43 proteinopathy"

### Extended Data Figures

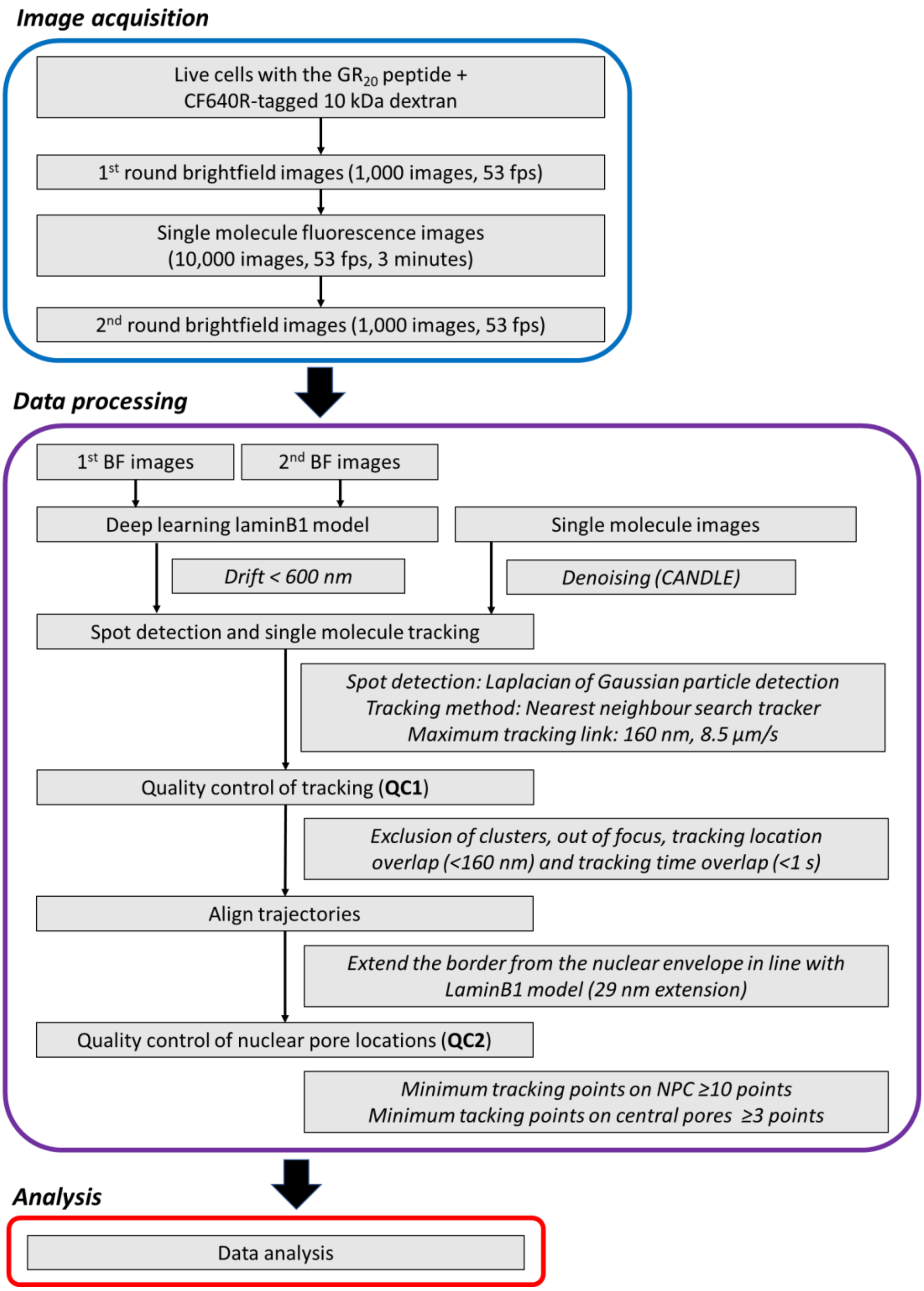


***Extended Data Figure 1.*** *Workflow of image acquisition, data processing and analysis.*

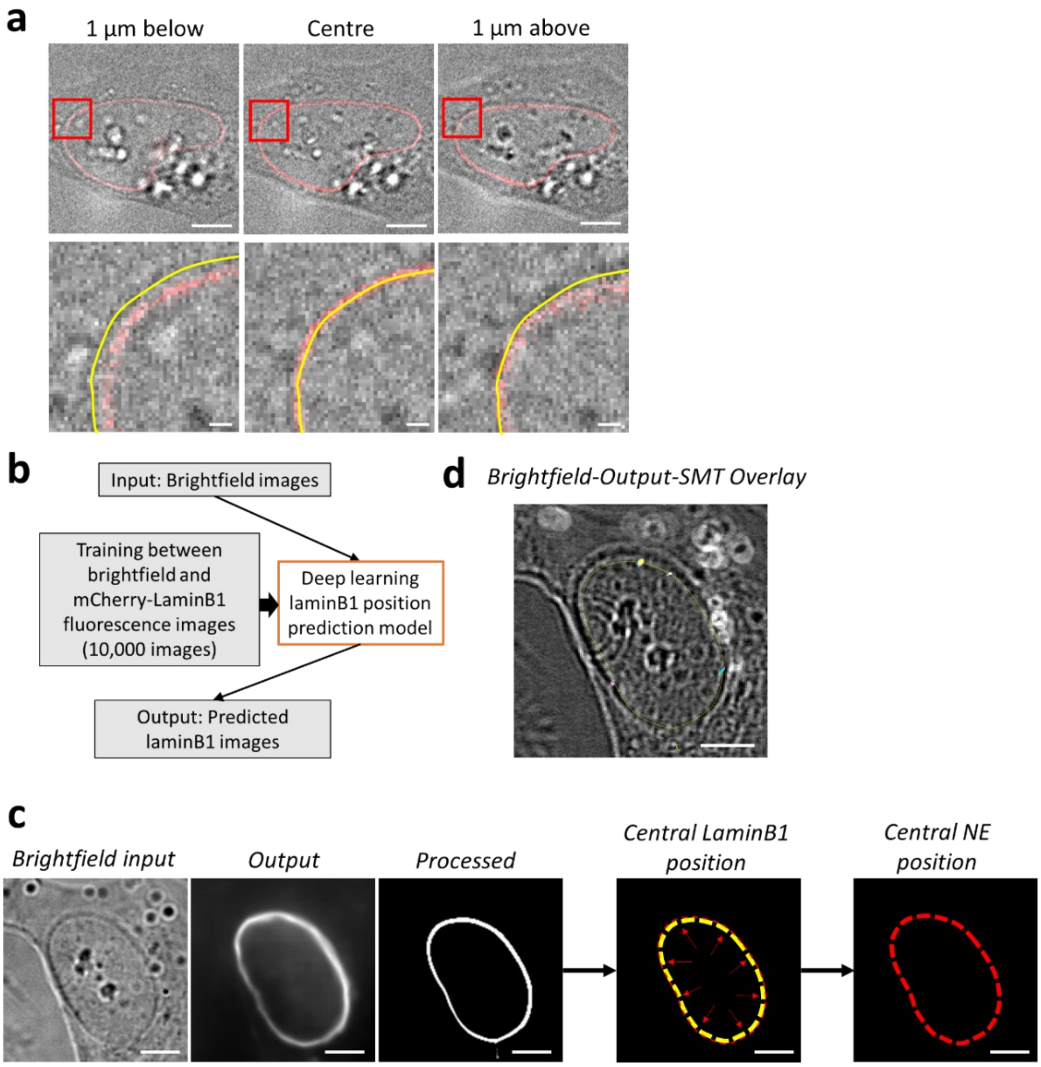


***Extended Data Figure 2. Deep learning analysis workflow for nuclear envelope positioning in non-labelled cells using brightfield imaging.*** *a) Combined brightfield and LaminB1 immunofluorescence images at different focal depths, with magnified views shown in red boxes in lower panels. Images from the central position of the nucleus show alignment between the midpoint of the nuclear envelope determined from brightfield (yellow line) and mCherry-LaminB1 fluorescence (red line) images, whereas images above and below the central focal plane show misalignment, showing need for correction of brightfield-based nuclear envelope positioning for single molecule tracking at the lower edge of the nucleus. b) Schematic representation of the deep learning process and model for nuclear envelope positioning using brightfield imaging. Training was performed on 10,000 images for brightfield and the nuclear envelope marker mCherry-lamin B1 to generate the deep learning laminB1 position prediction model. c) Example of deep learning image processing. Two sets of bright-field images (1,000 pre-tracking and 1,000 post-tracking) were averaged to generate the input representation. The deep learning model then generated the LaminB1 position prediction as the output. This output was further processed to enhance contrast and sharpen the boundary, improving accuracy of nuclear envelope's central position definition. The central position of the LaminB1 position was determined (yellow line) and then a 29nm outward correction was applied for definition of the nuclear envelope central position (red line) to account for the known Lamin B1 localisation to the inner nuclear membrane34, 35. d) Combined image showing single molecule tracking trajectories (multi-coloured lines) overlaid on the brightfield image with the nuclear envelope central position as determined by the deep learning model (yellow line). Scale bar is 5 µm for top panels in a) and c) and d), 1 µm for c) lower panels.*

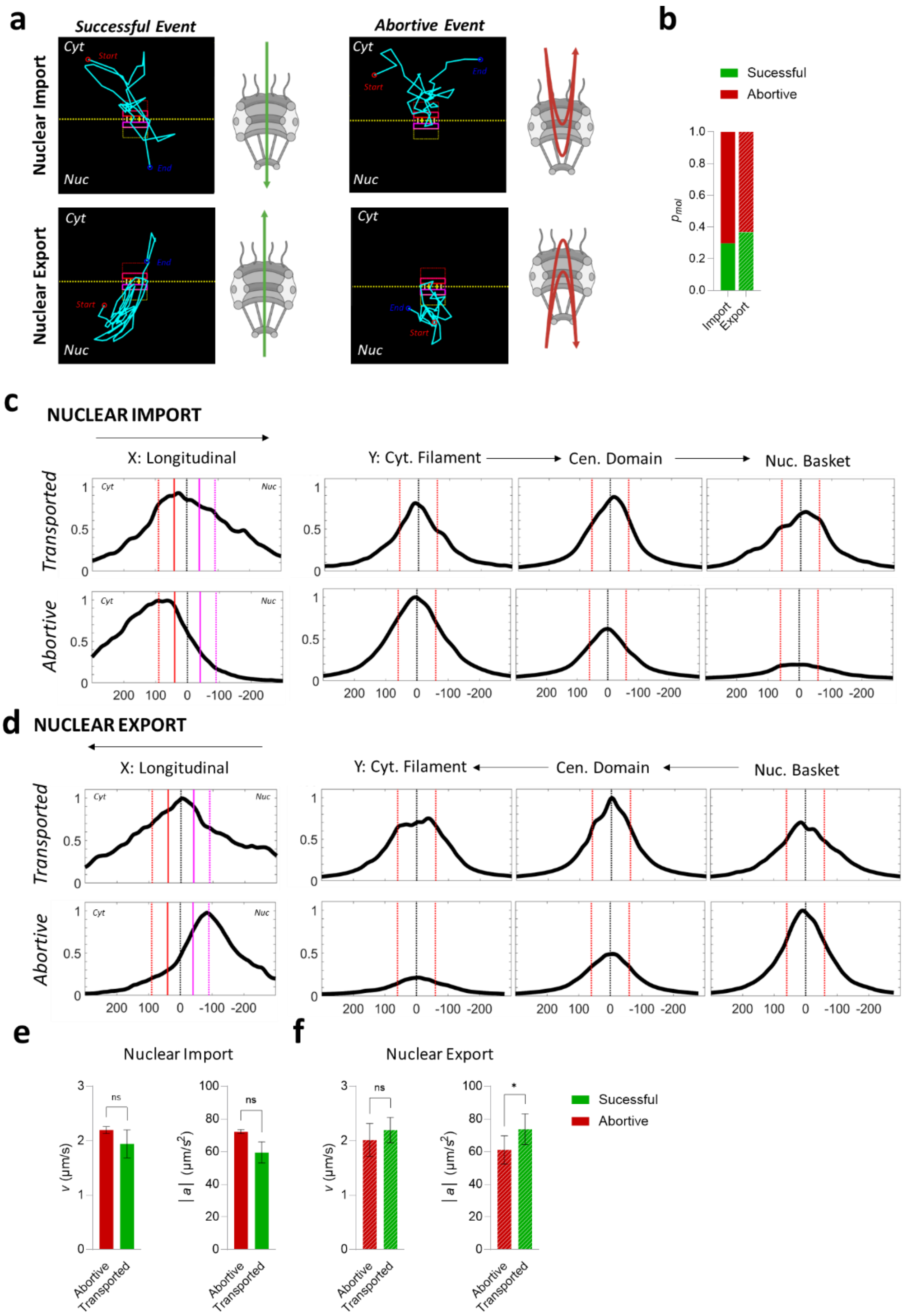

***Extended Data Figure 3. Single molecule nucleocytoplasmic transport dynamics comparing nuclear import and export, and successfully transported and abortive track dynamics.*** *a) NPC schematic and representative track examples of nuclear import and export events with successfully transported and abortive trajectories; scale bars are 100 nm. b) Comparative analysis of proportional of molecular tracks (pmol) during nuclear import and export (diagonal striped bars) for successfully transported (green) and abortive (red) trajectories. Comparative longitudinal density profile through the NPC and transverse profiles through each of the cytoplasmic filament, central and nuclear basket subdomains (as per Figure 1) for successfully transported and abortive trajectories during nuclear import (d) and nuclear export (e); arrows indicated direction of transport. Comparative analysis of mean velocity (v) and absolute acceleration (|a|) during nuclear import (solid bars, e) and export (diagonal striped bars, f) for successfully transported (green) and abortive (red) trajectories. Statistical analyses of experimental data: paired t-test on log-transformed data; *p<0.05; N=3; raw numerical data in Source Data File for Extended Data Figure 3.*

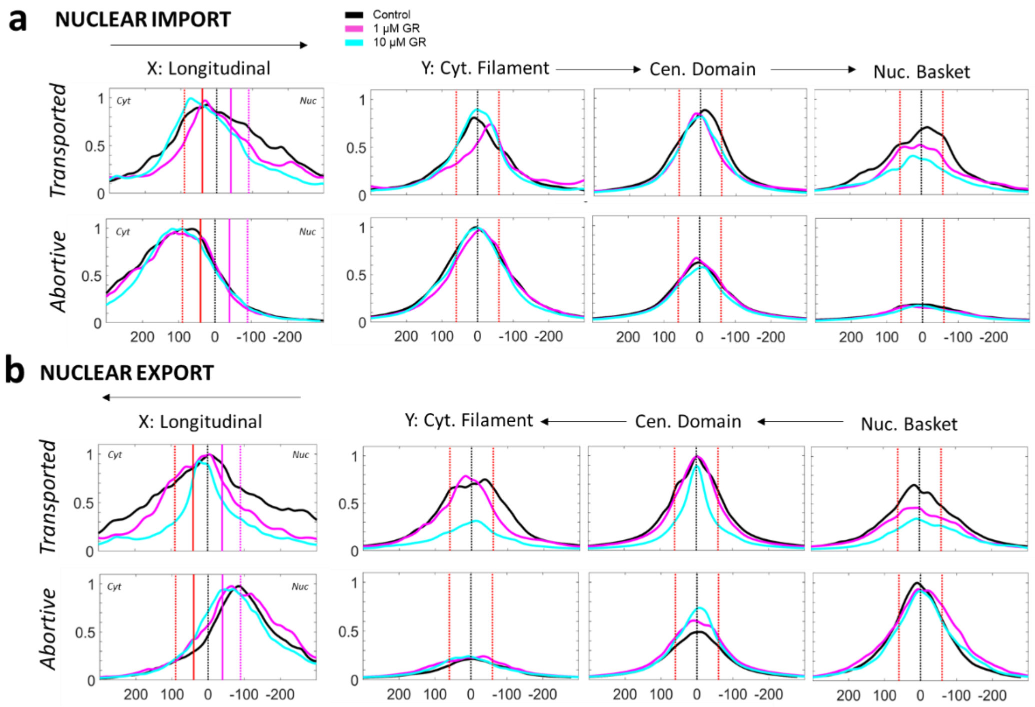


***Extended Data Figure 3. Single molecule nucleocytoplasmic transport dynamics upon polyGR treatment comparing successfully transported and abortive track dynamics for nuclear import and export.*** *Comparative longitudinal density profile through the NPC and transverse profiles through each of the cytoplasmic filament, central and nuclear basket subdomains (as per Figure 1, control – black, 1μM GR20 - magenta, 10μM GR20 - cyan) for successfully transported and abortive trajectories during nuclear import (a) and nuclear export (b); arrows indicated direction of transport.*

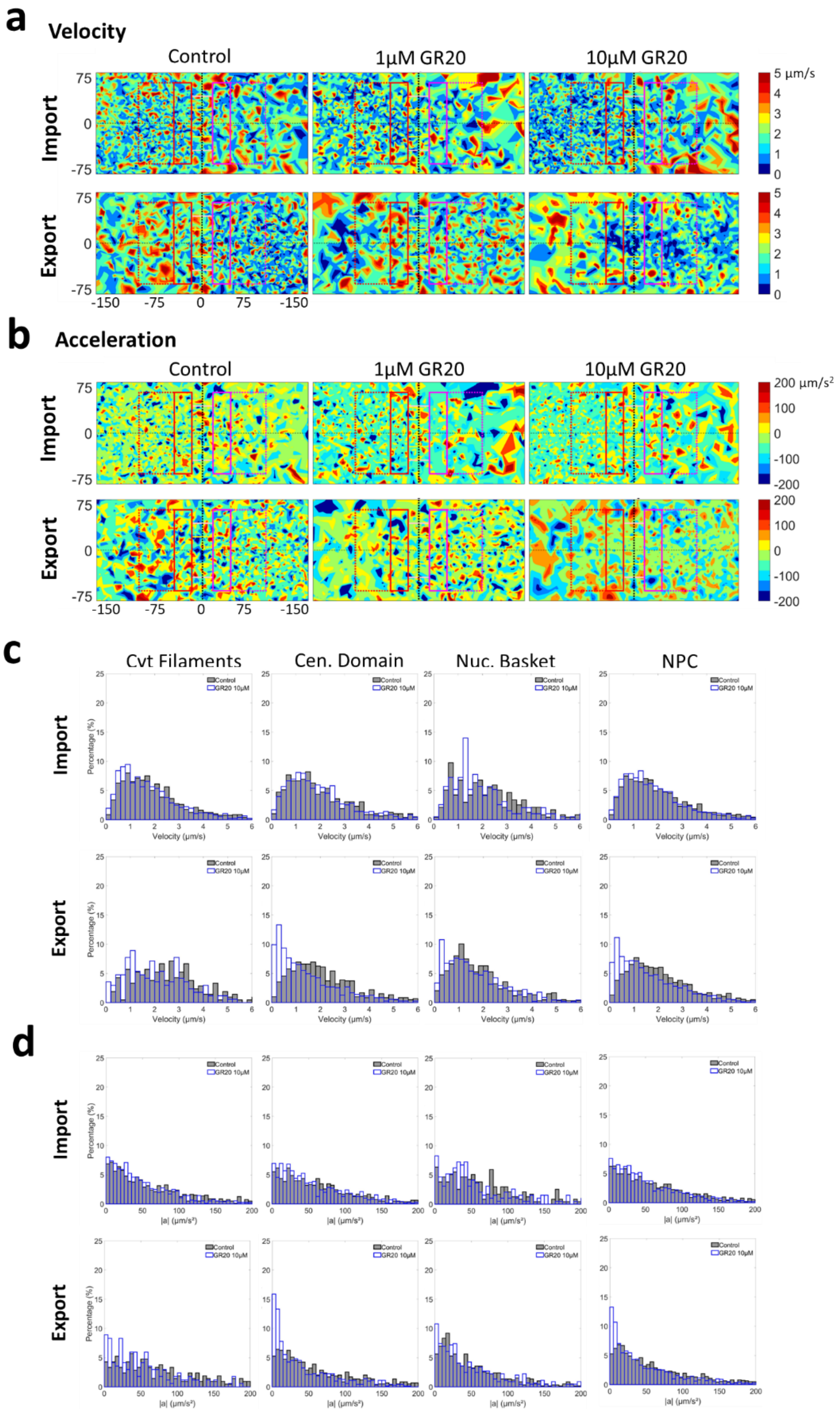


***Extended Data Figure 5. Density heatmaps and histograms for nucleocytoplasmic transport dynamics in untreated and polyGR peptide treated cells segregated into import and export trajectories as per data presented in Figure 2c and e.*** *Density heatmaps with superimposed NPC dimension model of (a) velocity and (b) absolute acceleration for import and export single molecule transport dynamics showing the variation across the NPC and subdomains; axis in nm Histograms of (c) velocity and (d) absolute acceleration for single molecule dynamic in the total NPC and cytoplasmic filament, central region, nuclear basket subdomains. Raw numerical data can be found in Source Data File for Figure 2.*

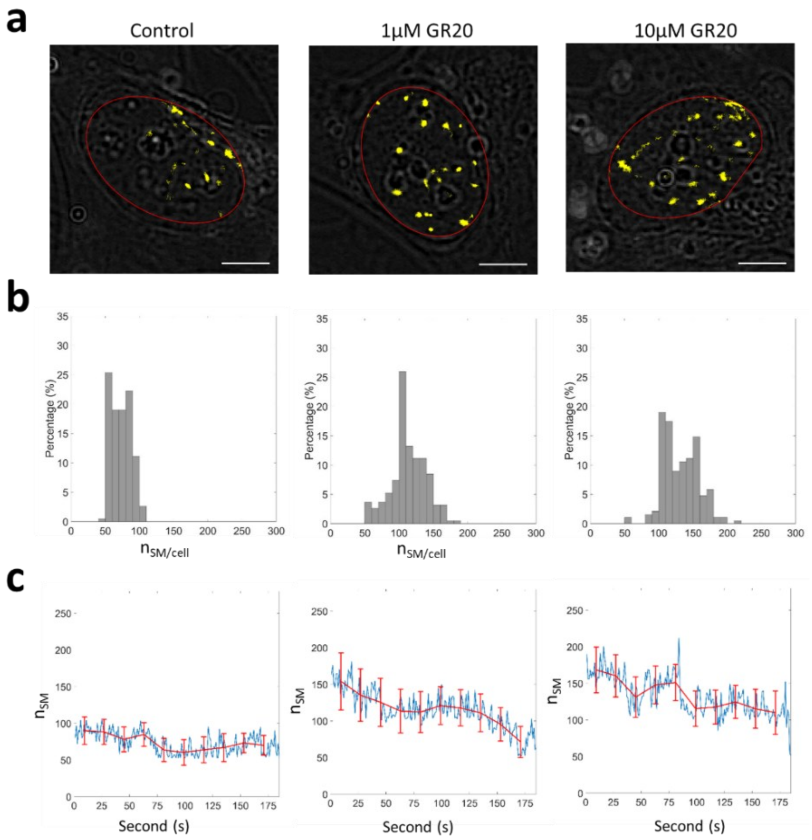


***Extended Data Figure 5. Cumulative single molecule nuclear accumulation and dynamics.*** *Data from single molecule tracking experiments were reanalysed to assess nuclear levels of 10 kDa dextran single molecules in untreated and polyGR (1 µM and 10 µM) treated cells. a) Example of brightfield images with single molecule detection (yellow) within the deep-learning defined nuclear envelope central position boundary (red) during the 3-minute experimental acquisition, scale bars are 5 µm. b) Histograms of average single molecule count per second per cell ($n_{SM/cell}$) showing that at a cumulative level, $GR_{20}$ treatment increases nuclear 10KDa dextran. c) Average single molecule count per second ($n_{SM}$) was plotted over the 3-minute experimental acquisition time-course with line fitting and SEM error (red), demonstrating a decrease over time with polyGR treatment that is not present in untreated cells. Raw numerical data can be found in Source Data File for Extended Data Figure 5.*

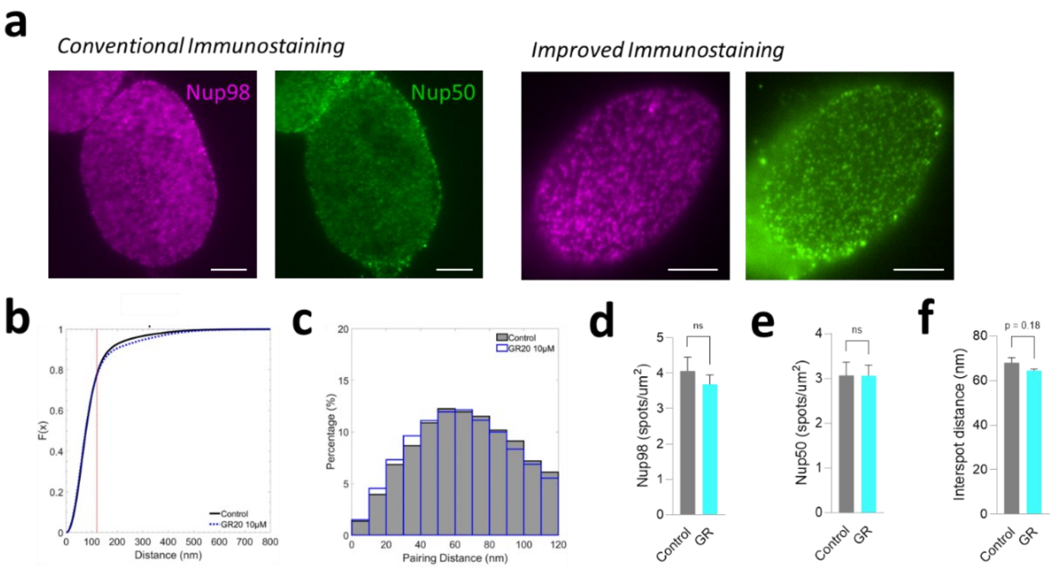


***Extended Data Figure 6. Further detail of paired localisation and diameter simulation analysis for NPC structure in relation to Figure 3.*** *a) Comparative fluorescent image acquisition from conventional and improved immunostaining methods, with clear improvement in definition of NPC spot localisations for Nup98 and Nup50; scale bars are 5μm. b) Cumulative distribution function of the measured inter-spot pairing distance between Nup98 and Nup50 for control and polyGR treated experimental data, with line (red) indicating 120 nm corresponding to the outer NPC diameter, after which pairing data is not considered. c) Histogram of Nup98 and Nup50 pairing distance within 120nm in control and polyGR-treated conditions. NPC spot detection density is measured for (d) Nup98 and (e) Nup50. f) Mean inter-spot distance between Nup98 and Nup50. Statistical analyses of experimental data: paired t-test; exact value given when p<0.2 to indicate trend; N=3; raw numerical data in Source Data File for Figure 3.*
